# A protein language model for domain, functional site, and variant effect prediction

**DOI:** 10.64898/2026.11.03.764008

**Authors:** Peng Wang, Mian Umair Ahsan, Tingting Chen, Kai Wang

**Affiliations:** Center for Cellular and Molecular Therapeutics, Children’s Hospital of Philadelphia, Philadelphia, PA 19104, USA; Bioengineering Graduate Group, University of Pennsylvania, Philadelphia, PA 19104, USA; Department of Pathology and Laboratory Medicine, University of Pennsylvania, Philadelphia, PA 19104, USA

## Abstract

Protein sequence databases are growing far faster than experimentally curated annotations, limiting functional interpretation of newly identified proteins and coding variants. We developed PALM, a protein annotation language model that jointly predicts domain and functional-site occurrence, while preserving overlapping and nested annotations through a soft-target objective. PALM integrates ESM2 with low-rank adaptation (LoRA), learned layer aggregation, and a bidirectional long short-term memory (BiLSTM) encoder. PALM was trained on ∼15 million InterPro-annotated UniRef50 sequences spanning 17,951 domain and 1,383 site classes. Across 2.3 million held-out proteins, PALM achieved domain- and site-occurrence F1 scores of 0.941 and 0.877, respectively, and retained substantial performance in proteins with <20% sequence identity to the training and validation data. PALM also substantially outperformed ESM3 annotation baseline. Without training on variant-pathogenicity labels, PALM embedding disruption distinguished 151,823 ClinVar missense variants more accurately than ESM2 embedding disruption (AUROC 0.835 versus 0.710), and combining PALM embedding disruption with the ESM2 likelihood-ratio signal reached an AUROC of 0.877. The same combined zero-shot score outperformed AlphaMissense on a benchmark dataset on 128,161 missense variants, that were derived from literature but not present in ClinVar database. At the protein level, PALM embeddings showed stronger CATH-Gene3D superfamily organization than frozen ESM2 and ESM3 embeddings, and improved prediction of experimental activation and repression effects when combined with ESM2. In a downstream case study, we demonstrated the ability to use PALM-derived embedding as proxy of deep mutational scanning (DMS), showcasing the broad application of PALM. Together, PALM converts curated residue-level knowledge into transferable and mechanistically interpretable representations that connect protein sequence, function, and variant consequence.

## Introduction

Advances in sequencing have made protein discovery routine, but expert curation has not kept pace with sequence growth. For example, UniProt release 2026_01 contains 203,130,941 protein entries, whereas only 574,627 are reviewed Swiss-Prot records, illustrating the scale of the annotation gap ^1^. To alleviate this problem, UniRef reduces redundancy for scalable sequence analysis, and InterPro integrates signatures from multiple member databases into a unified vocabulary of protein families, domains, repeats, and functional sites ^2, 3^. The same interpretive gap affects human genetics: ClinVar continues to accumulate missense submissions, many of which lack definitive clinical interpretation ^4^. For instance, among all ClinVar variant records (∼1.1 millions) added in 2025, 61.75% of them remain classified as “uncertain significance” on May 3, 2026.

Sequence-alignment and profile-based methods for protein sequence annotation remain foundational. BLAST identifies local similarity to characterized sequences, whereas profile hidden Markov models (HMM) can detect more remote family relationships ^5, 6^. These approaches are accurate and biologically interpretable when suitable reference families exist, but large-scale annotation can require extensive database searches and post-processing. Moreover, a protein may contain nested or overlapping domains and several site types at the same residue, creating a structured multi-label problem that is not fully represented by a single annotation label. For example, an active site could sit on top of a long transmembrane domain, correctly predicting both the site and domain will be important for functional interpretation as well as building a more meaningful representation of protein.

Protein language models (pLMs), such as ESM2, offer a complementary route by learning contextual residue representations from large collections of natural sequences ^7, 8^. ESM2 ^8^ was trained using millions of nature-occurring protein sequences from UniRef ^3^, and model learnt to predict the corrupted (masked) amino acids based on the rest of the protein sequences. By learning from millions of wild-type protein sequence, model inherently learnt biochemical properties and evolutionary conserveness of proteins. Their embeddings capture structural and functional information, and masked-marginal likelihood scores can prioritize functional or disease-associated substitutions without task-specific training ^9, 10^. There are other masked-language-models trained on both wild-type and mutant protein sequences to enhance the variant effect prediction, such as MutFormer ^11^. However, masked-language-model pretraining is optimized to recover amino-acid distributions, not to emit explicit domain and site labels. Functional knowledge and interpretability encoded in curated annotation resources such as InterPro ^2^ therefore remains only indirectly represented in those models.

Deep learning has also been applied directly to protein annotation. These methods usually used sequence and annotation from Pfam ^12^, which uses curated protein sequence instead of nature-occurring sequence and has been integrated into InterPro ^2^. ProtCNN used neural networks to classify sequence by Pfam family and ProtENN took majority vote across an ensemble of 19 ProtCNN models to reduce error rates ^13^. In addition, there are transfer-learning studies that evaluated pLM representations for functional annotation under limited labeled data ^14^. PSALM uses ESM2 representations, a residue-level domain-state classifier, and a structured probabilistic decoder to produce coherent domain calls ^15^. These advances establish the value of learned representations, but a single non-overlapping domain segmentation does not address the joint prediction of overlapping domain and site annotations. Moreover, limited variant-effect interpretation and downstream generalizability have been shown in these methods, underestimating the potential of annotation-enhanced protein language model.

To address these limitations, we developed PALM, a Protein Annotation Language Model that learns residue-level domains and functional sites from naturally occurring UniRef sequences and InterPro annotations. PALM integrates ESM2 with low-rank adaptation (LoRA), learned layer aggregation, and a bidirectional long short-term memory (BiLSTM) encoder. Domain and site annotations are modeled with separate occurrence and classification heads that share a unified residue representation, while a soft-target objective allows each residue to retain multiple valid labels and therefore represent overlapping annotations. Because all annotation tasks are learned through the same representation, PALM also produces functionally informed embeddings that can transfer beyond annotation prediction. We evaluate PALM on held-out proteins across sequence-similarity strata, compare its annotation performance with ESM3 ^16^, and examine downstream representation quality using CATH-Gene3D superfamilies ^17^, Reactome pathways ^18^, ClinVar variants ^4^, PubMind variants ^19^, and FlexiNR mutational screens results ^20^. By unifying overlap-aware protein annotation, transferable representation learning, and interpretable mutation-disruption signals, PALM provides a framework for connecting protein sequence to function and variant consequence.

## Results

### Construction of an InterPro-annotated protein dataset

We constructed a large residue-annotated dataset by mapping InterPro records to UniRef50 representative sequences ^2, 3^ (**Supplementary Figure 1**). InterPro identifiers and coordinate ranges were stored in FASTA headers, enabling compact on-disk storage and deterministic label reconstruction during training. The resulting dataset contained 15,365,469 proteins: 11,521,114 for training, 1,534,749 for validation, and 2,309,606 in the held-out test set (**Table 1**). The annotation vocabulary comprised 17,951 domain classes and 1,383 site classes.

**Table 1.** Composition of the InterPro-annotated UniRef50 dataset and the sequence-identity evaluation subset. For better evaluate the performance in held-out dataset, we stratified the test dataset into different groups, based on the protein sequence identity (PID, %) comparing with train + validation datasets (T+V).

| Dataset component | Definition | Number of Protein Sequences | Dataset proportion (% of full dataset) |
|---|---|---|---|
| Training | Model fitting | 11,521,114 | 74.98% |
| Validation | Checkpoint selection and monitoring | 1,534,749 | 9.99% |
| Held-out test | Final annotation evaluation | 2,309,606 | 15.03% |
| Total dataset | InterPro-annotated UniRef50 representatives | 15,365,469 | 100.00% |
| Train + validation (T+V) | Deduplicated reference sequence for sequence similarity calculation | 13,055,863 | 84.97% |
| Test subset stratified by PID comparing to T+V | Subset of held-out test after PID calculation and stratification | 686,370 | 4.47%<br>(29.72% of held-out test set) |

To assess performance across sequence similarity, we generated multiple test subsets each with different sequence similarity (range from 0% to 100%) compared to the training and validation datasets. By calculating protein sequence identity (PID) in held-out test set compared to train + validation sets, the test subsets together contained 686,370 test proteins, corresponding to 29.72% of the full test set (**Supplementary Table 1**).

To characterize InterPro annotations across UniRef50 sequences, we examined protein length and annotation frequency, interval length, and sequence coverage (**Table 2**). Site annotations included functional, active, conserved, repeat, and post-translational modification sites. Site annotations were substantially sparser than domain annotations: they occurred in only 15% of proteins, compared with 97% for domains, and covered an average of 4% of each sequence in terms of total length, compared with 24% for domains. Site intervals are also shorter, averaging 18.56 amino acids versus 104.09 amino acids for domains (in train set).

**Table 2.** The statistics of domain and site annotation in different datasets. The annotation (i.e., InterPro ID) on a protein could have duplication because same domain or site could occur in two discontinuous regions on one protein.

| Data Split | Annotation type | Average protein length (aa) | Average annotation length (aa) | Average number of annotation per protein | Percentage of protein with at least one annotation | Average length coverage of annotation |
|---|---|---|---|---|---|---|
| Train | Domain | 438.86 | 104.09 | 3.20 | 97.08% | 23.72% |
|  | Site | 438.86 | 18.56 | 0.21 | 14.95% | 4.23% |
| Validation | Domain | 428.02 | 104.77 | 3.14 | 97.02% | 24.48% |
|  | Site | 428.02 | 18.53 | 0.21 | 15.13% | 4.33% |
| Test | Domain | 422.87 | 104.25 | 3.12 | 97.06% | 24.65% |
|  | Site | 422.87 | 18.48 | 0.20 | 14.94% | 4.37% |

### Model design and overlap-aware training

**Figure 1** summarizes how PALM connects residue-level annotation learning with downstream protein and variant analysis. By learning domain and site occurrence and identity from a shared representation, PALM captures multiple functional features at the same residue, including overlapping annotations (**Figure 1a**). The same model is then applied to wild-type and mutant sequences, allowing changes in annotations and embeddings to quantify potential functional disruption (**Figure 1b**). This design enables us to test whether annotation supervision supports not only accurate protein annotation but also zero-shot variant-effect prediction and broader functional representation (**Figure 1c**).

**Figure 1.**
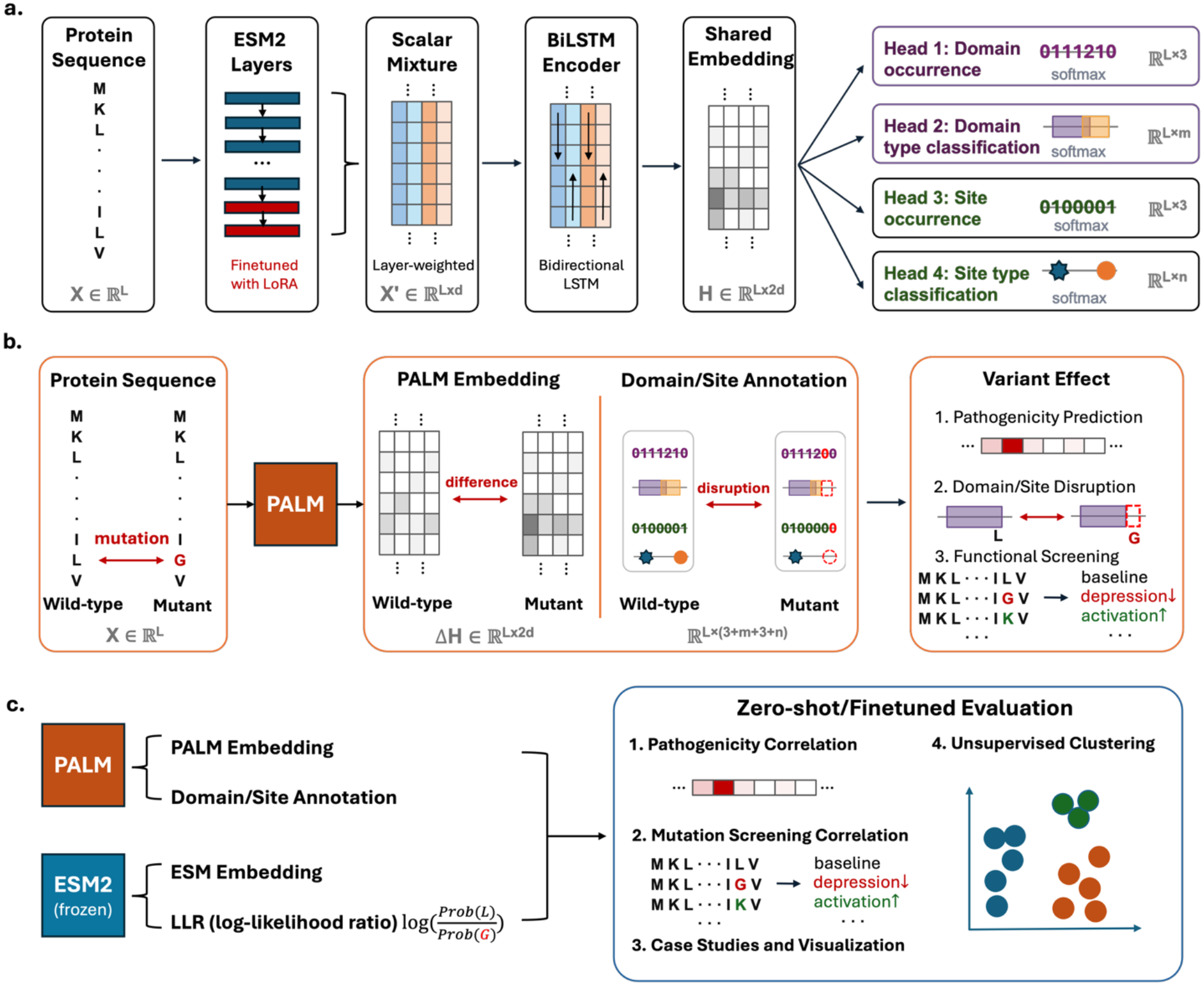
PALM architecture and evaluation design. **(a)** Protein sequences are encoded by ESM2-150M, whose 30 transformer layers produce 640-dimensional residue states. LoRA adapters update selected modules, and a learned scalar mixture combines layer representations before a bidirectional LSTM and four prediction heads. **(b)** Domain and site occurrence heads predict no, single, or multiple annotations; class heads predict InterPro identities with soft-assignment. **(c)** The model is evaluated on held-out annotation data and downstream variant- and function-prediction tasks.

To select an initial ESM model and downstream model structures for use in this study, we compared different parameter sizes of ESM2 model, tested finetune approaches such as LoRA adaptation, and evaluated different downstream encoders including Transformer, BiLSTM, and multilayer perceptron (**Supplementary Table 2**). Based on these pilot experiments, we selected ESM2-150M as the protein-sequence encoder and a BiLSTM as the final encoder before the prediction heads. We then evaluated four strategies for residue-level prediction: conventional multiclass classification, independent multilabel classification, a detection transformer (DETR) model, and a mask-transformer model (**Supplementary Figure 2**). After selecting the multiclass framework (Strategy 1) as baseline, we compared alternative training objectives and adopted a soft-target loss that preserves overlapping protein annotations (Strategy 1b and 1c, **Supplementary Figure 3**). Traditional method collapsed the domain and site annotation, forcing model to predict only one final classification per residue, but PALM resolved this limitation by applying a soft-target loss during model training (**Figure 2**). The final PALM design used shared sequence representations with four task-specific heads: domain occurrence, site occurrence, domain class, and site class (**Figure 2b**). The occurrence heads distinguished residues with no, one, or multiple annotations, whereas the class heads assigned InterPro labels. This factorization separated whether and how many annotations occur from which labels are present.

**Figure 2.**
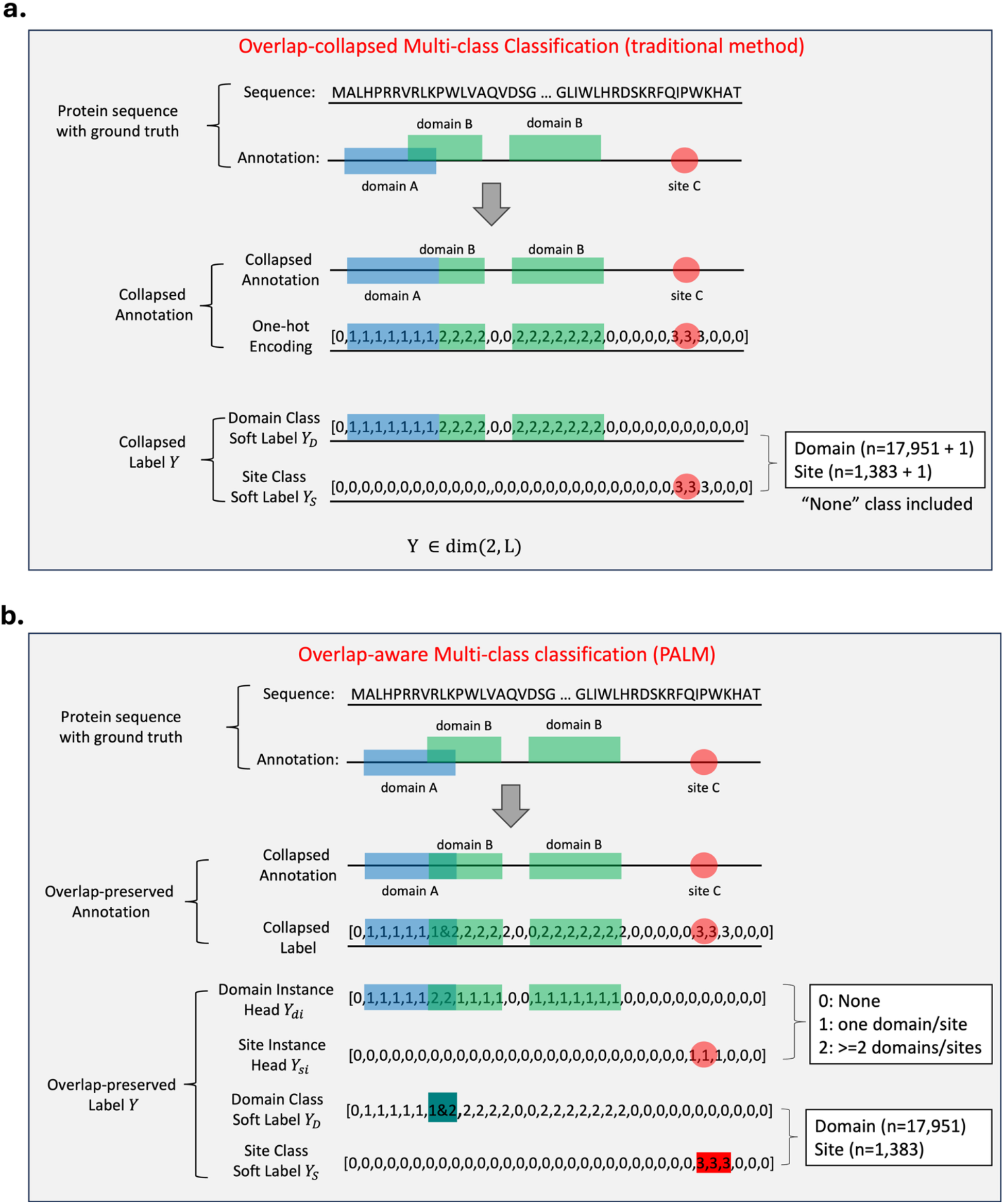
The comparison of traditional domain classification and the soft-target classification in PALM. **(a)** Traditional domain and site classification treats each residue as a multiclass classification problem and retains only one final domain or site class. **(b)** PALM uses a soft-target multiclass objective. InterPro annotations are collapsed while retaining overlapping labels, and target probability is distributed across all valid domain or site classes. This objective encourages recovery of multiple annotations when they overlap at a residue.

For residues with overlapping labels, the class objective distributed target mass equally across all valid classes. Unlike a hard single-label target, this soft-target formulation penalized a model that concentrated all probability on only one member of a valid label set. In development experiments, this objective trained more reliably than the set-prediction alternatives while preserving the ability to represent overlapping domain and site annotations.

### PALM accurately predicts residue-level domain and site annotations

On all 2,309,606 held-out proteins, the soft-target PALM model achieved domain-occurrence precision of 94.49%, recall of 93.77%, and F1 of 94.13%. Functional sites are substantially sparser than domains (Table 2) and are more challenging to predict, yet PALM still achieved good site-occurrence precision at 88.37%, recall at 87.12%, and F1 at 87.74% (Table 3). Relaxed top-class accuracy was 97.14% for domain type and 99.45% for site type. In the supported-label comparison shown in Figure 3, PALM also substantially exceeded ESM3 across residue-level occurrence and classification tasks. These results indicate strong residue-level performance for both annotation types while highlighting the greater difficulty of sparse and heterogeneous site annotations.

**Figure 3.**
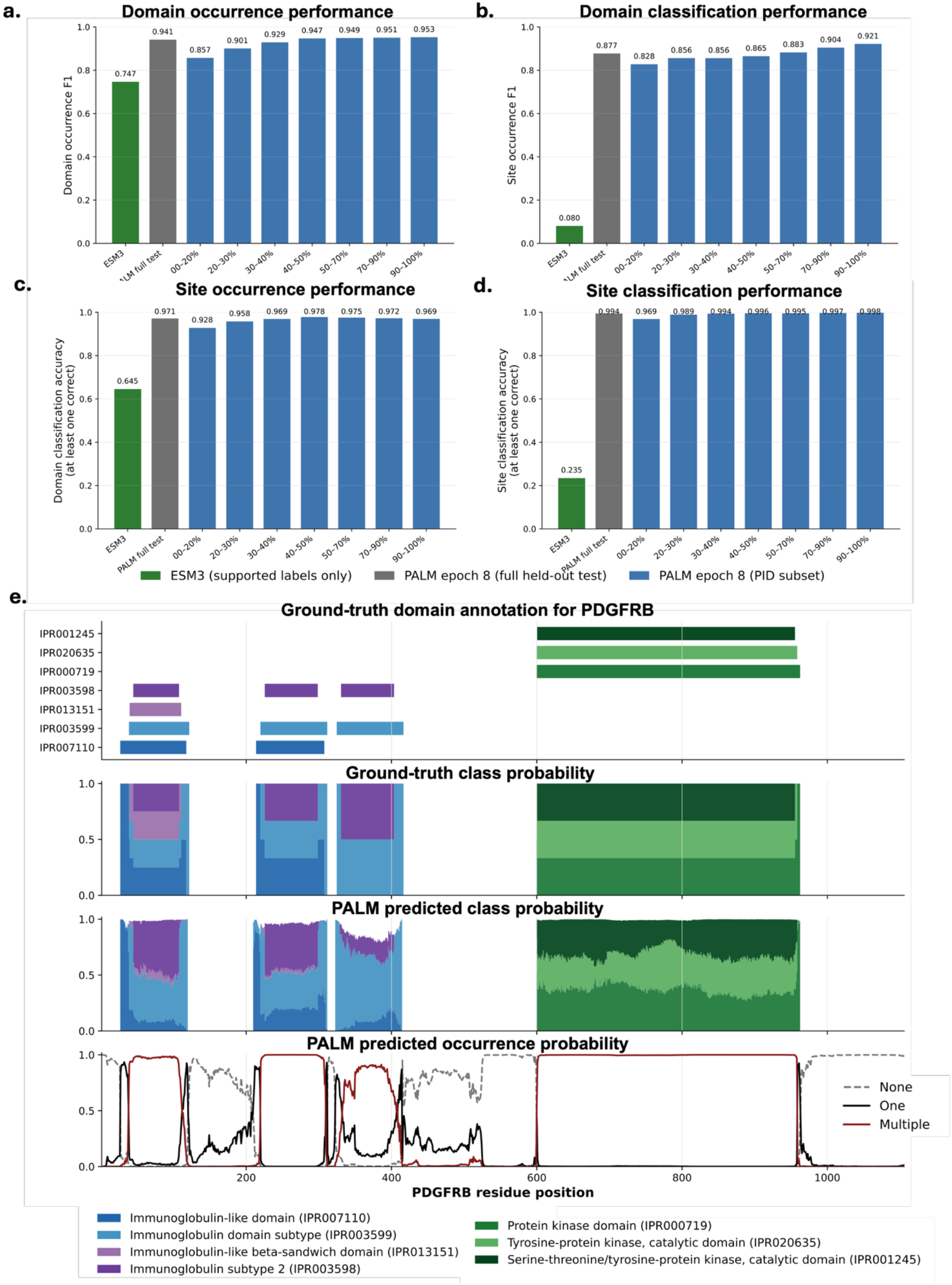
Residue-level annotation performance and an example prediction. (a-d) PALM performance across the full held-out set and the completed PID-stratified subset for domain occurrence, site occurrence, domain classification, and site classification; ESM3 function-annotation performance on supported labels is shown as an external comparison. (e) PALM predictions for platelet-derived growth factor receptor beta (PDGFRB), showing ground-truth domain annotation, ground-truth derived class probability, PALM predicted domain probability, and PALM predicted domain occurrence probability.

**Table 3.** Protein domain and site annotation performance in the held-out test dataset (n=2,309,606). The precision*, recall*, F1* for occurrence heads are calculated after collapsing three instance classes into binary prediction: none as 0, single or multiple domains as 1.

| **Model head** | **Prediction task** | **Number of classes** | **Precision\*** | **Recall\*** | **F1\*** | **Accuracy** |
|---|---|---|---|---|---|---|
| Domain occurrence | none, single, or multiple domains | 3 | 94.49% | 93.77% | 94.13% | 93.17% |
| Site occurrence | none, single, or multiple sites | 3 | 88.37% | 87.12% | 87.74% | 99.78% |
| Domain-type | InterPro domain classification | 17,951 | / | / | / | 97.14% |
| Site-type | InterPro site classification | 1,383 | / | / | / | 99.45% |

Performance remained robust across the PID-stratified subset (**Figure 3a-d**). PALM achieved domain- and site-occurrence F1 scores of 0.941 and 0.877, respectively, across the completed 686,370-protein subset. In the lowest PID bin, containing proteins with less than 20% maximum sequence identity to the training-plus-validation reference or no detectable hit, the corresponding F1 scores were 0.857 and 0.828. Performance increased with similarity to the training reference; the descriptive sum of the two occurrence F1 values increased from 1.685 in the lowest bin to 1.874 in the highest bin. This trend is consistent with sequence-related generalization, while the low-PID results show that predictions were not restricted to near-duplicate proteins.

To illustrate PALM’s ability to predict overlapping annotations, we selected PDGFRB, a multidomain receptor tyrosine kinase containing five extracellular immunoglobulin-like domains, a transmembrane region, an intracellular kinase domain, and regulatory sites that mediate downstream signaling ^21^. PALM recovered most of the domain annotations, including multiple ground truth labels at overlapping residues (**Figure 3e**), demonstrating the importance of comprehensive multilabel prediction for capturing the protein’s layered functional architecture.

### PALM provides complementary and interpretable zero-shot signals for variant pathogenicity

Having established PALM’s ability to annotate domains and functional sites in wild-type reference proteins, we next asked whether these learned annotations and representations could inform missense variant-effect prediction. dbNSFP ^22^ catalogs many established predictors with strong performance on ClinVar variants, whereas PALM was trained only on UniRef protein sequences and InterPro annotations; neither ClinVar nor any mutation database was used during PALM training. We therefore evaluated PALM in a fully zero-shot setting on 151,823 ClinVar missense variants classified as pathogenic, likely pathogenic, benign, or likely benign ^4^.

Because functional sites are sparse and heterogeneous (Table 2), we did not assume that simply falling within a predicted site is sufficient to determine pathogenicity. Instead, we tested whether mutation-induced changes in PALM domain/site predictions and learned embeddings provided complementary variant-effect signals. To reduce dependence on ClinVar, we also evaluated PALM and external predictors on PubMind ^19^, which uses large language models to extract variant–disease–pathogenicity evidence from literature abstracts and full text and has only ∼10% overlap with ClinVar. We analyzed coding variant records across all PubMind confidence levels (n=158,213) and a higher-confidence subset with confidence ≥2 (n=97,610).

On ClinVar, the ESM2 masked-marginal likelihood ratio (LLR) achieved an AUROC of 0.844. Combining the normalized LLR with PALM annotation disruption modestly increased the AUROC to 0.850, whereas combining it with PALM embedding disruption increased the AUROC to 0.877 (**Figure 4a**). PALM embedding disruption alone achieved an AUROC of 0.835 and a PR-AUC of 0.705, substantially exceeding the corresponding ESM2 embedding-difference scores (AUROC, 0.710; PR-AUC, 0.510). These results indicate that annotation training reorganized the protein representation space to capture variant effects that were not fully reflected by the frozen ESM2 embeddings.

**Figure 4.**
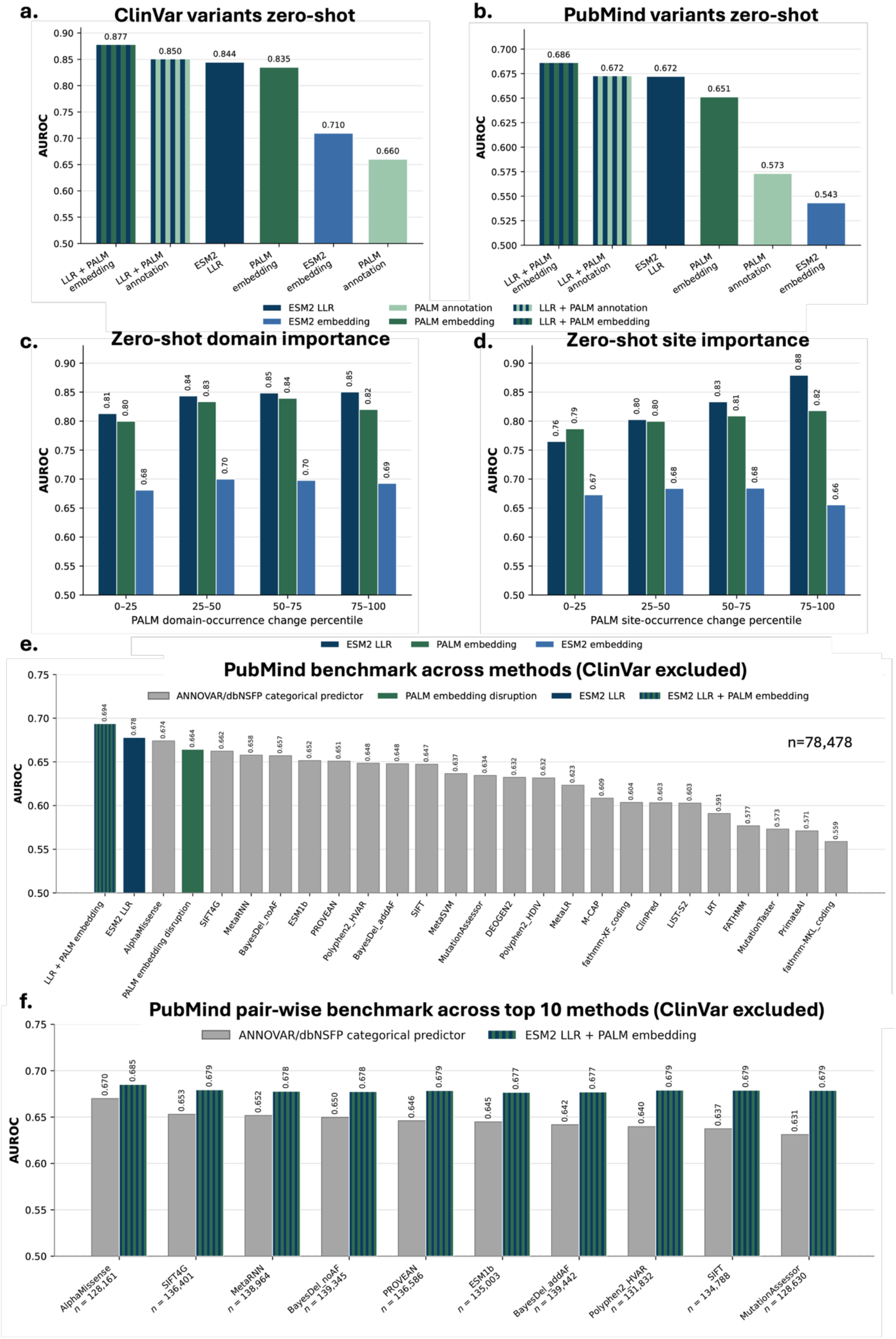
Zero-shot pathogenicity prediction using the final PALM model. The final soft-target, overlap-aware PALM and frozen ESM2-150M were applied to wild-type and mutant protein sequences without training on ClinVar or PubMind pathogenicity labels. **(a)** AUROC on 151,823 binary-labeled ClinVar missense variants (48,042 pathogenic/likely pathogenic and 103,781 benign/likely benign). **(b)** Record-level AUROC on 97,610 PubMind records with confidence ≥2 (89,444 pathogenic and 8,166 benign). ESM2 LLR is the masked-marginal log-likelihood ratio, log P(WT) - log P(mutant). PALM annotation and embedding scores quantify mutation-induced disruption of the domain/site prediction heads and learned protein representations, respectively.

The complementary signal generalized to PubMind. ESM2 LLR achieved an AUROC of 0.672, PALM embedding disruption achieved 0.651, and their equal-weight combination achieved the highest AUROC of 0.686 (**Figure 4b**). PALM outputs additionally provided residue-level context for interpreting variant effects. When ClinVar variants were stratified by predicted site-occurrence disruption, ESM2 LLR performance increased from an AUROC of 0.765 in the lowest quartile to 0.879 in the highest quartile (**Figure 4d**). A similar, although smaller, increase was observed across domain-occurrence disruption quartiles (**Figure 4c**). Thus,

PALM provides both a complementary zero-shot pathogenicity score and an interpretable measure of the functional context in which a substitution occurs.

In order to provide a fair comparison of PALM’s zero-shot abiality with other softwares and tools that are likely to train with ClinVar already. We used PubMind variants, which are derived from biomedical literatures directly and have been shown with low overlap with ClinVar ^19^. On the ClinVar-exclusive complete-case PubMind benchmark across all confidence levels, the combined zero-shot score also achieved the highest AUROC (0.694), compared with 0.674 for AlphaMissense and 0.678 for ESM2 LLR (**Figure 4e**). Pairwise comparisons between the combined zero-shot score and the ten highest-ranked categorical tools likewise showed consistently better AUROC in either ClinVar-exclusive PubMind variants (Figure 4f) or ClinVar-inclusive PubMind variants (Supplementary Figure 4a).

To evaluate PALM’s standalone performance in variant pathogenicity prediction without ESM2 LLR, we fine-tuned PALM on ClinVar variants and evaluated it on the ClinVar-exclusive PubMind benchmark (Supplementary Figure 4b,c). PALM achieved AUROCs of 0.658 and 0.659 under random and gene/protein-disjoint splits, respectively, ranking 2^nd^ and 3^rd^, slightly below AlphaMissense (0.665). Both PALM models achieved the highest PR-AUCs among the evaluated predictors (0.887 and 0.888, respectively), with comparable performance across the two splitting strategies.

Overall, we showed that PALM embedding showed strong ability for variant pathogenicity prediction, and the performance could be further boosted when combined with untouched ESM2 frozen embedding. The combined score outperformed all other methods in PubMind benchmark even in the zero-shot setting. By finetuning using ClinVar variants, PALM standalone achieved best balanced performance in PubMind variants with AUROC and PR-AUC are both used.

Combined scores are equal-weight means of cohort-standardized ESM2 LLR and PALM scores. **c, d,** ClinVar AUROC stratified by quartiles of the Jensen–Shannon divergence between wild-type and mutant PALM predictions for domain occurrence (c) or site occurrence (d) at the mutated residue. **e,** Record-level comparison with 23 dbNSFP v4.7 categorical predictors on a strict complete-case subset of 78,478 ClinVar-exclusive PubMind records across all confidence levels (65,750 pathogenic and 12,728 benign). Categorical calls were collapsed to a consensus call, and every displayed method was evaluated on the same records. Bars are ordered by AUROC. **f**, Pairwise comparison of the combined ESM2 LLR and PALM embedding-disruption score with the top 10 categorical predictors. Each pair uses its own matched non-ClinVar subset,the total number of variants in each pairwise comparison is shown in the plot.

### PALM embeddings more accurately capture protein superfamily relationships

To examine whether residue-level domain and site supervision improves the organization of protein-level representations, we evaluated PALM embeddings using CATH-Gene3D annotations ^17^, which assign protein domains to homologous superfamilies based on structural classification and sequence homology. The benchmark comprised 5,449 reviewed canonical human proteins spanning 62 CATH-Gene3D superfamilies. We compared mean-pooled embeddings from frozen ESM2-150M, ESM3-sm-open-v1, and PALM checkpoints using leave-one-out cosine k-nearest-neighbor (kNN) classification at k = 5, 10, and 20. Each protein was excluded from its own neighbor search and assigned a superfamily based on the majority label among its k nearest neighbors (Figure 5a).

**Figure 5.**
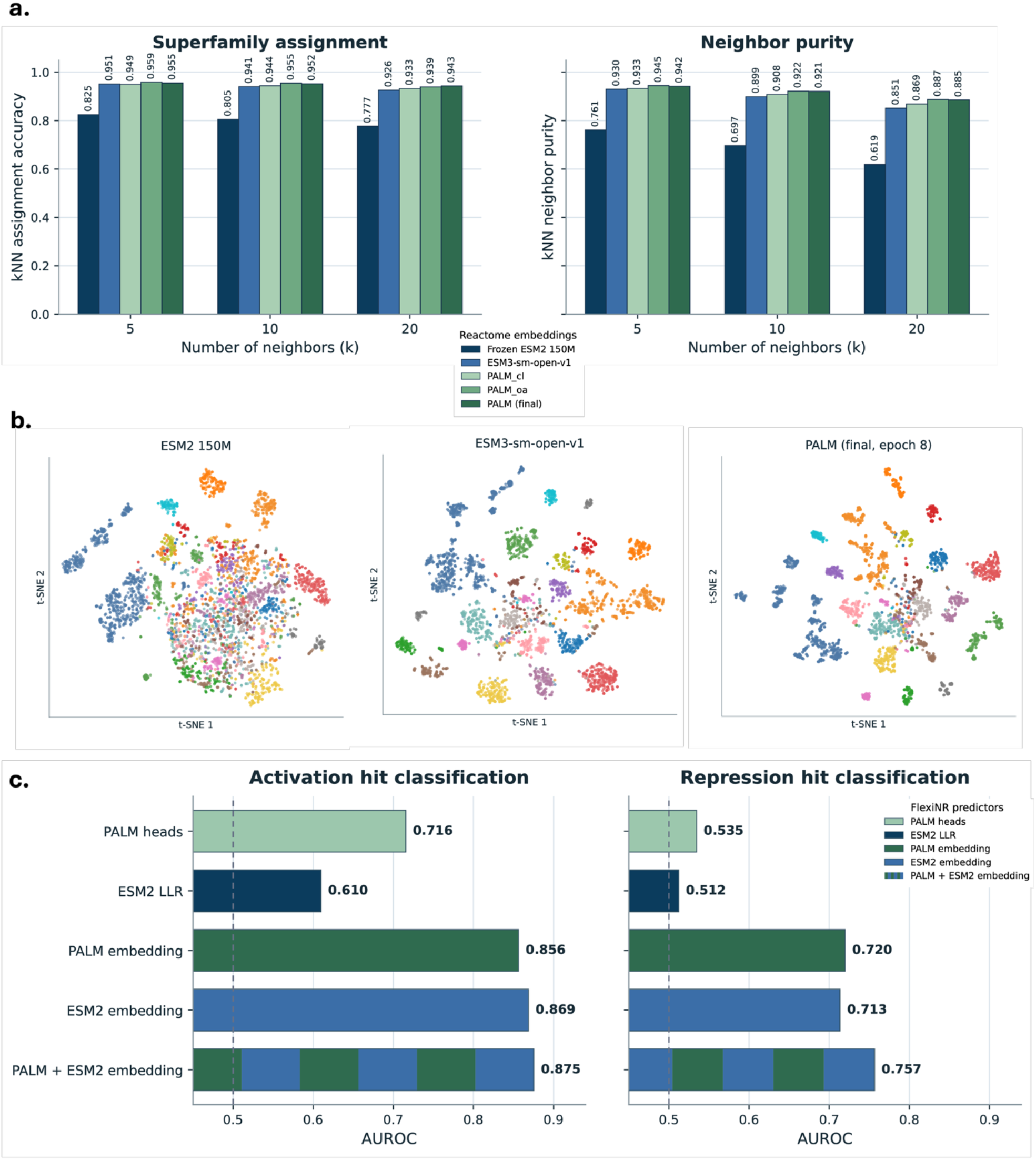
PALM representations capture protein function and variant effects. **(a)** CATH-Gene3D superfamily assignment accuracy and neighbor purity for frozen ESM2-150M, ESM3-sm-open-v1, and PALM embeddings across k = 5-20 nearest neighbors. **(b)** t-SNE visualization of ESM2, ESM3, and final PALM embeddings for proteins from 20 representative CATH-Gene3D superfamilies; PALM shows clearer separation and more compact superfamily-associated groups. **(c)** Five-fold group-cross-validated AUROC for predicting changes in FlexiNR activation and repression hit status (450 wild-type-mutant pairs per screen) using PALM heads, ESM2 LLRs, individual embeddings, or concatenated PALM-ESM2 embeddings. Dashed lines indicate chance performance.

PALM produced stronger superfamily organization than the pretrained ESM baselines. At k = 5, assignment accuracy was 0.955 for the final PALM model, compared with 0.951 for ESM3 and 0.825 for ESM2; neighbor purity was 0.942 for PALM, 0.930 for ESM3, and 0.761 for ESM2 (Figure 5a). The same ordering was maintained across the tested neighborhood sizes, indicating that annotation supervision reshapes the protein representation beyond the local residue-level prediction tasks.

We further visualized the embedding space using t-SNE plot for proteins from 20 representative CATH-Gene3D superfamilies (Figure 5b). ESM2 embeddings showed substantial overlap among superfamilies, whereas ESM3 separated several groups more clearly. PALM produced the most distinct and compact superfamily-associated clusters overall, consistent with the kNN analysis. Together, these results suggest that residue-level domain and site supervision improves organization of PALM embeddings at the protein-superfamily level.

Because CATH-Gene3D superfamily annotations are related to the InterPro annotations used to train PALM and to functional annotations available to ESM3, this benchmark is not fully independent. We therefore retained Reactome pathway analysis as a complementary benchmark in Supplementary Figure 5. Across 11,312 human proteins assigned to 16 top-level Reactome labels, PALM embeddings showed higher kNN pathway-assignment accuracy and neighbor purity than frozen ESM2, with ESM3 included as an additional embedding baseline in the updated visualization. This complementary analysis supports the conclusion that PALM’s representation captures broader functional organization beyond the CATH-Gene3D benchmark.

Taken together, the CATH-Gene3D and Reactome analyses indicate that domain and site supervision improves the organization of protein embeddings at both homologous-superfamily and broader pathway levels, while the overlap between CATH-Gene3D and InterPro annotations warrants conservative interpretation of the superfamily benchmark.

### PALM embeddings provide complementary signals for mutational-screen prediction

We further evaluated whether PALM representations capture functional effects measured by FlexiNR deep mutational screens ^20, 23^. Using the native 80-residue assay sequences, we classified whether perturbations changed hit status in the activation and repression screens (450 wild-type–mutant pairs per screen), using five-fold group cross-validation by tile identity. Predictors used changes in PALM annotation-head outputs, ESM2 log-likelihood ratios (LLRs), PALM or ESM2 embeddings, or concatenated PALM–ESM2 embeddings.

The concatenated representation achieved the highest AUROC in both screens: 0.875 for activation, compared with 0.869 for ESM2 and 0.856 for PALM embeddings alone, and 0.757 for repression, compared with 0.713 and 0.720, respectively (Figure 5c). Embedding-based predictors also substantially outperformed PALM annotation heads (0.716 and 0.535) and ESM2 LLRs (0.610 and 0.512). These results indicate that PALM and ESM2 embeddings encode complementary functional information, with their integration providing a particularly clear improvement for repression-hit classification.

### PALM provides localized, interpretable readouts of variant effects

Most pathogenicity predictors summarize a variant with a single score, making it difficult to determine why a prediction conflicts with the observed phenotype. We use the PTEN C124F variant to illustrate this challenge. The variant affects the catalytic Cys124 residue and receives high pathogenicity scores from several computational predictors, whereas a literature-derived PubMind record (PubMind access ID: PVID762957) links C124F to a reported case with benign thyroid histopathology. This represents a phenotype-prediction discordance rather than evidence that C124F itself is benign. PALM provided additional local resolution: the predicted active-site occurrence signal was largely preserved for C124F, whereas the pathogenic C124S control produced a more pronounced reduction around the catalytic region (**Figure 6 a-f**).

**Figure 6.**
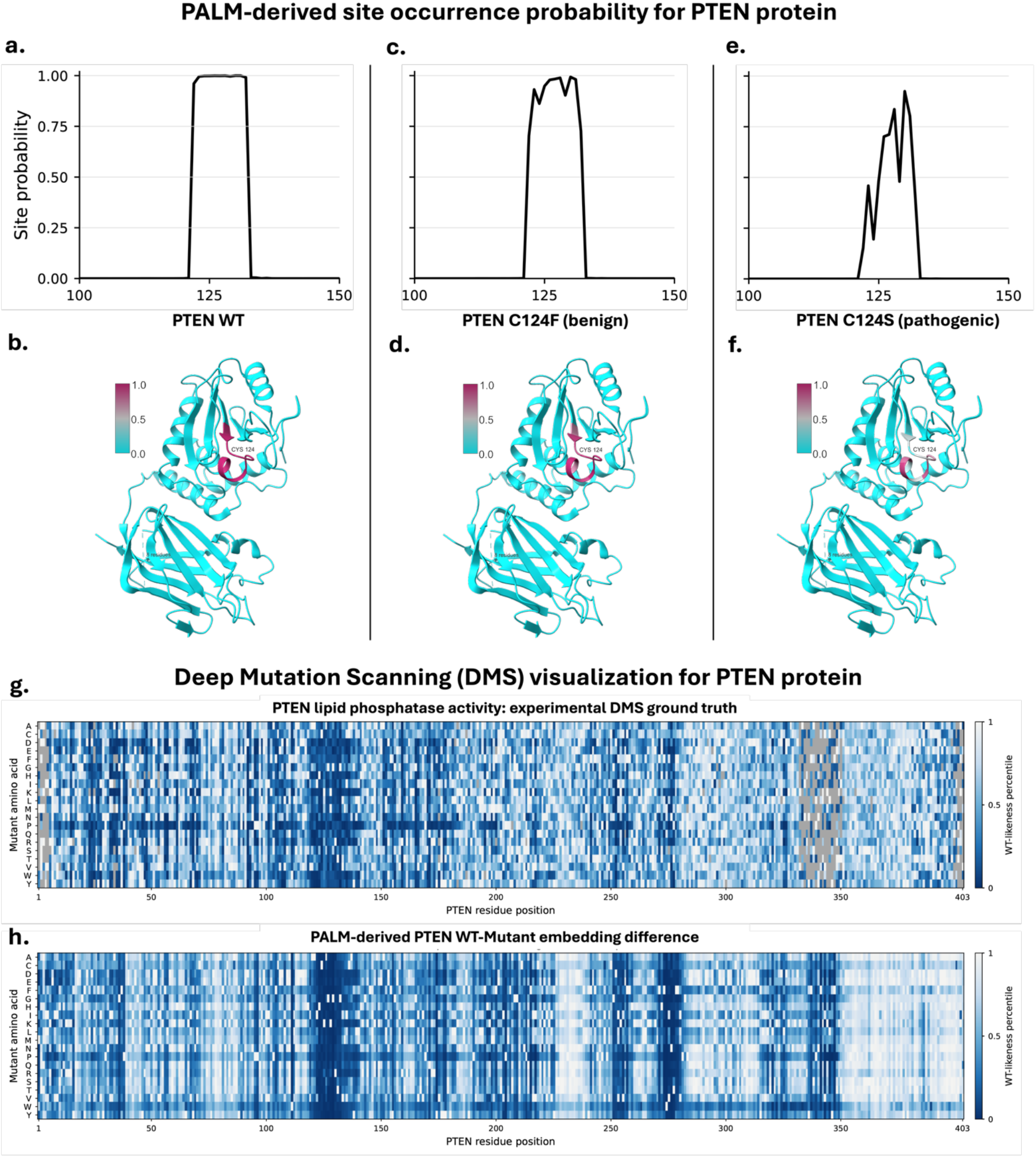
PALM annotation readout for PTEN variant-effect interpretation. PALM site-occurrence probability is shown along the sequence and mapped onto the PTEN X-ray structure (PDB 1D5R; residues 7-353) for wild-type PTEN (**a, b**), the PubMind-linked benign C124F variant (**c, d**), and pathogenic C124S (**e, f**). Site-occurrence probability is the summed probability of the single-site and multiple-site occurrence states. The predicted Cys124 active-site signal is largely preserved for C124F but more strongly disrupted by C124S. **g**, Experimental deep mutational scanning (DMS) measurements of PTEN lipid phosphatase activity, displayed as variant fitness percentiles; dark grey indicates missing DMS data. **h**, PALM-derived WT–mutant embedding differences across PTEN missense substitutions. In both heatmaps, columns represent residue positions and rows represent substituted amino acids, enabling comparison of experimentally measured functional effects with PALM-derived embedding changes.

To examine whether PALM representations also reflect experimentally measured functional effects, we visualized PTEN lipid phosphatase activity from deep mutational scanning (DMS) ^24^ alongside PALM-derived wild-type–mutant embedding differences (Figure 6g, h). Across 7,260 missense variants, PALM embedding-derived WT-likeness correlated with experimental PTEN lipid phosphatase activity (Spearman *ρ* = 0.426; Pearson *r* = 0.484). After averaging substitutions by residue position, the association increased (Spearman *ρ* = 0.597; Pearson *r* = 0.580; n = 403 positions). The heatmaps showed qualitative regional correspondence, with reduced experimental fitness and greater embedding changes concentrated in several shared regions, including the catalytic region. Together, these readouts provide complementary views of predicted annotation changes and broader representation changes. Preservation of an active-site annotation signal and zero-shot embedding difference do not establish preserved enzymatic activity or clinical benignity, yet they provide proxy and new functional interpretation of human genetic variants.

## Discussion

In this study, we developed PALM, a protein annotation language model that simultaneously predicts residue-level domains and functional sites while preserving overlapping annotations. PALM achieved strong annotation performance across more than 2.3 million held-out proteins, retained substantial performance in the lowest sequence-identity subset, and substantially exceeded the supported-label ESM3 ^16^ annotation baseline shown in Figure 3. Domain and site supervision also produced functionally informed embeddings that organized CATH-Gene3D superfamilies ^17^ more clearly than frozen ESM2 ^8^ and ESM3 ^16^ embeddings, while Reactome ^18^ provided a complementary pathway-level benchmark. These representations further supplied complementary zero-shot signals for variant-effect prediction across ClinVar ^4^ and PubMind ^19^ and improved experimental mutational-screen prediction like FlexiNR ^20^ when combined with ESM2. Together, these findings show that explicit residue-level annotation learning can extend protein language models beyond sequence likelihood toward scalable and interpretable connections among protein sequence, function, and variant consequence.

PALM addresses the gap between rapidly expanding protein-sequence collections and comparatively limited residue-level functional annotation. Homology-based methods such as BLAST and profile HMMs remain indispensable for interpreting proteins with suitable reference families ^5, 6, 25^, whereas PALM provides a complementary, single-pass framework that predicts domains and functional sites at every residue.

The central methodological contribution is the treatment of annotation overlap. PSALM , for example, combines ESM2 features with a residue classifier and structured probabilistic decoder to produce a single coherent domain segmentation ^15^. PALM instead separates annotation occurrence from class identity and predicts domains and sites in parallel. Unlike a non-overlapping segmentation, this design accommodates nested domains, coincident site types, and multiple valid labels at the same residue. The soft-target objective further rewards probability assigned across all valid annotations rather than accepting only one correct member of an overlapping label set.

The downstream benchmarks suggest that annotation supervision changes the information encoded by the protein representation. In ClinVar, PALM embedding disruption markedly outperformed ESM2 embedding disruption and complemented ESM2 LLR; the same combination also provided the strongest zero-shot performance in PubMind. In CATH-Gene3D, PALM produced higher kNN superfamily assignment and neighbor purity than ESM2 and ESM3, with UMAP showing clearer superfamily-associated structure (Figure 5a&b). Reactome pathway organization and FlexiNR mutational-screen prediction provided complementary evidence that PALM embeddings capture functional relationships not fully represented by sequence likelihood or ESM2 embeddings alone. Importantly, the CATH-Gene3D result should not be interpreted as a fully independent benchmark because CATH-Gene3D and InterPro annotations are related; its value is strongest when considered alongside the more independent Reactome and experimental-screen evaluations (Supplementary Figure 5).

PALM’s residue-level outputs provide a distinct interpretability benefit. A scalar pathogenicity score can be considered together with the predicted domain or site at the affected residue and the mutation-induced change in that annotation. In the PTEN example, PALM distinguished the predicted local active-site consequences of C124F and C124S despite both substitutions affecting the same catalytic residue, and the structure-based visualization made the localized disruption directly inspectable (Figure 6). Such differences can generate mechanistic hypotheses and help prioritize discordant cases for review. Moreover, PALM-derived WT–mutant embedding difference sharing regional correspondence with PTEN lipid phosphatase activity from deep mutational scanning (DMS) ^24^, indicating a broad application of using PALM as a proxy for whole protein scanning. Nevertheless, PALM outputs remain predictions learned from database annotations and only serve as proxy in the case study, but PALM still provide complementary interpretation when combining with other methods.

Our study has several limitations. First, InterPro integrates both curated knowledge and computationally inferred signatures ^2^. Therefore, incomplete annotations, correlated classes, and biases from individual member databases may propagate into PALM. In addition, some InterPro annotations are primarily derived from sequence similarity rather than direct experimental evidence of protein function, contributing to heterogeneity in domain and site labels. Second, there is currently no standardized benchmark for comprehensive residue-level protein annotation. We therefore evaluated PALM directly on held-out InterPro proteins and sequence-identity-stratified subsets, while recognizing that InterPro annotations do not always provide a complete representation of protein function. To complement this direct evaluation, we further assessed the learned representations using CATH-Gene3D superfamily assignments ^17^ and Reactome pathway annotations ^18^. Nevertheless, more comprehensive and standardized benchmarks will be needed to enable rigorous comparison among protein annotation methods. Third, ClinVar and PubMind were used to evaluate downstream variant-effect prediction, but both resources have inherent limitations. ClinVar is a submission-based database with variable coverage and evidence quality, whereas PubMind is an LLM-derived literature resource and may inherit uncertainties from information extraction, variant normalization, and inconsistencies across published studies ^4, 19^. Experimental deep mutational scanning (DMS) datasets, such as the FlexiNR activation and repression screens ^20^ evaluated here, partially address these limitations by providing systematic functional measurements across many substitutions within a protein. However, most DMS experiments remain focused on individual proteins or functional assays and therefore provide limited coverage of the broader proteome. Despite these constraints, PALM achieved strong domain and functional-site annotation performance, outperformed ESM3 ^16^ in the evaluated annotation setting, and addressed overlapping residue annotations through a soft-target learning objective. Importantly, the downstream analyses suggest that preserving this overlapping functional information is not only useful for annotation itself, but also produces representations that better capture protein relationships and variant effects.

Future work should incorporate InterPro evidence quality and source-specific confidence during training, improve representation of rare annotation classes, and calibrate model probabilities to provide more reliable confidence estimates. For variant interpretation, PALM and future protein annotators should be evaluated on experimentally characterized DMS datasets spanning a broader range of proteins, functional mechanisms, and variant types, including insertions, deletions, and truncating variants. Incorporating structural context, evolutionary conservation, and molecular interaction information, such as protein–protein interactions, may further improve predictive performance while retaining the interpretability of annotation-based outputs. Notably, our results also show that combining evolution-derived ESM2 representations with annotation-informed PALM representations improves several downstream predictions, suggesting that these representations capture complementary aspects of protein biology. Future protein models may therefore benefit from integrating multiple complementary modalities, including sequence evolution, functional annotation, interaction profiles, and three-dimensional structural representations, rather than relying on sequence information alone. Within these limitations, PALM provides a scalable framework for translating protein sequences into residue-level functional annotations and annotation-aware variant representations, while providing a foundation for multimodal models that more directly connect sequence variation to protein function and disease mechanisms.

## Materials and Methods

### Data sources and preprocessing

UniRef50 representative sequences were used as the sequence source ^3^ (release Date “2025-10-08”; version “2025_04”; URL: http://uniprot.org/uniref). InterPro records (version 1.0) were mapped to the representative sequences and divided into domain and site vocabularies ^2^. Only proteins with at least one mapped InterPro annotation were retained. Each FASTA header stored a compact string containing InterPro identifiers and residue-coordinate ranges; residue-level labels were reconstructed from these intervals during training and evaluation.

The domain vocabulary contained 17,951 classes, and the site vocabulary contained 1,383 classes, including functional sites, repeats, conserved sites, and post-translational modification (PTM) sites. Domain and site targets were constructed independently, allowing a residue to carry both types and multiple labels of the same type. Overlap-aware targets were stored as sparse label sets to avoid materializing dense tensors over the complete vocabulary.

### Dataset splitting

The 15,365,469 proteins were partitioned on disk into 11,521,114 training, 1,534,749 validation, and 2,309,606 held-out test proteins. FASTA files contained approximately 10,000 records, yielding 1,162 training files, 155 validation files, and 233 test files. The validation set was used for checkpoint selection and training monitoring; the held-out test set was reserved for final annotation evaluation.

### PID-stratified test grouping

To estimate generalization as a function of sequence similarity, exact duplicate sequences were removed from the combined training and validation reference by sequence SHA-1 hash. The resulting reference contained 13,055,863 proteins. Held-out proteins were searched against this reference with BLASTP ^6, 25^, retaining up to five candidate hits per query. The best hit was selected by query-adjusted protein sequence identity (PID), followed by bit score and then local percent identity.

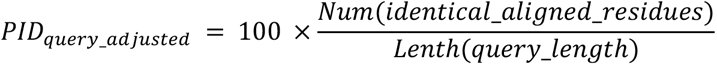

PID bins were [0,20), [20,30), [30,40), [40,50), [50,70), [70,90), and [90,100]. Coverage classes used an 80% threshold for both query and target coverage. The full BLASTP search was expensive; consequently, PID-stratified evaluation used the leading completed 686,370 test proteins (29.72% of the full held-out split).

### Model architecture

PALM used the 150-million-parameter ESM2 model (ESM2-150M) as its backbone ^8^. Its 30 transformer layers produce 640-dimensional residue representations. Backbone parameters were frozen except for LoRA adapters applied to selected transformer modules in the final layers ^26^. Hidden states were combined by a learned scalar mixture ^27^ and passed to a bidirectional LSTM ^28^.

The shared representation fed four task-specific multilayer perceptron heads. Domain- and site-occurrence heads predicted three states per residue: no annotation, one annotation, or multiple annotations. The domain-class head predicted 17,951 InterPro domain labels, and the site-class head predicted 1,383 site labels. Occurrence heads were trained on all residues; each class head was trained at residues containing the corresponding annotation type.

### Training objective

Occurrence heads used categorical cross-entropy over the three multiplicity states. For an annotated residue i with a set Si of valid labels, the soft-target class loss was:

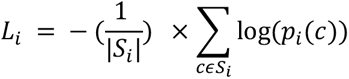

where *p_i_*(*c*) is the class-head probability assigned to label c. The total training objective was a weighted sum of the domain-occurrence, site-occurrence, domain-class, and site-class losses. The soft-target loss was compared with an earlier relaxed overlap-aware objective:

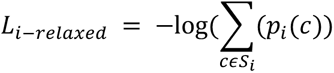

The relaxed objective is satisfied when probability concentrates on any one valid class. By averaging the log probabilities over all valid labels, the soft-target objective penalizes omission of other overlapping annotations.

### Annotation evaluation

Annotation performance was evaluated at residue level. Domain and site occurrence were converted to binary presence or absence and summarized by precision, recall, and F1 from pooled true-positive, false-positive, and false-negative counts. Class output was evaluated with relaxed top-class accuracy: a prediction was counted as correct when its highest-scoring class was present in the residue’s true label set. Metrics were computed on the full held-out test set and within each PID bin of the completed PID-stratified subset.

### ClinVar zero-shot variant scoring

The zero-shot analysis used 151,823 scorable ClinVar missense variants with pathogenicity classification benign, likely benign, pathogenic, or likely pathogenic ^4^. Pathogenic and likely pathogenic variants were contrasted with benign and likely benign variants; variants without one of these endpoint labels were excluded. For ESM2, the masked-marginal score was the log probability of the wild-type residue minus that of the mutant residue at the masked position ^9, 10^, so larger LLR values indicated greater predicted deleteriousness.

PALM was run separately on wild-type and mutant sequences. Domain-occurrence, site-occurrence, domain-class, site-class, aggregate annotation, and embedding-difference features were computed from the resulting output changes. Probability shifts and distribution-divergence summaries were oriented so that larger values represented greater disruption. Scores were standardized by z-score or percentile transformation before combination; the ESM2+PALM score used equal weights after normalization.

### ClinVar supervised fine-tuning

Downstream pathogenicity heads were trained using PALM-derived wild-type and mutant features. In the random-split design, variants were assigned randomly to training and test partitions. In the gene/protein-disjoint design, all variants from a gene/protein were restricted to one partition to reduce leakage from protein-specific patterns. Evaluation metrics were AUROC, average precision, accuracy, balanced accuracy, precision, recall, specificity, and F1.

### PubMind and external predictor evaluation

PubMind ^29^ was evaluated at record and coordinate-consensus levels for scorable missense variants with coordinate and transcript mappings. Scores included ESM2 LLR, PALM annotation disruption, and their normalized combination. ANNOVAR ^30^ and dbNSFP v4 ^22^ supplied external predictor values where available. Because predictor coverage differed, each comparison retained its own available sample size and was interpreted as a covered-subset analysis.

### CATH-Gene3D and Reactome embedding benchmarks

Protein-level embeddings were obtained by mean-pooling residue representations from PALM, frozen ESM2-150M ^8^, and ESM3-sm-open-v1 ^16^. For the primary representation benchmark, proteins were evaluated using CATH-Gene3D superfamily ^17^ assignments with leave-one-out cosine k-nearest-neighbor classification at k = 5, 10, and 20; assignment accuracy and neighbor purity were reported, and UMAP was used to visualize 20 representative superfamilies. Because CATH-Gene3D ^17^ and InterPro ^2^ contain related protein-family annotations, Reactome ^18^ top-level pathway labels were retained as a complementary benchmark on 11,312 human proteins. Reactome kNN analysis used k = 5, 10, 20, and 50 and reported pathway-assignment accuracy and neighbor purity, with UMAP used for qualitative comparison of embedding-space organization.

### Deep mutational scanning benchmarks

We evaluated PALM-derived variant-effect readouts using mutational-screen datasets measuring viral-effector activity ^20^ and PTEN lipid phosphatase activity ^24^. FlexiNR activation and repression measurements were obtained from the published viral-effector mutational screen ^23^ and analyzed in the context of the corresponding high-throughput protocol ^20^. Features included ESM2 LLR, PALM annotation-head disruption, PALM embedding differences, ESM2 embedding differences, and concatenated PALM+ESM2 differences. Both 80-residue local windows and full/cropped protein contexts were evaluated. Activation- and repression-hit classification was measured by AUROC; continuous effect-size prediction was measured by Spearman correlation.

PTEN DMS measurements were obtained from a published massively parallel functional assay that measured lipid phosphatase activity in a humanized yeast model ^24^, downloaded from MaveDB ^31^. The study reported high-confidence functional measurements for 7,244 single-amino-acid variants. For the PTEN visualization, we compared experimental scores for missense substitutions with whole-protein embedding differences between wild-type and mutant PTEN generated in zero-shot manner by PALM. Experimental scores and embedding differences were independently converted to percentile ranks scaled from 0 to 1 and displayed as matched heatmaps, with residue positions as columns and substituted amino acids as rows. Missing experimental measurements were shown in dark grey. This comparison enabled qualitative assessment of regional correspondence between experimentally measured functional effects and PALM-derived representation changes.

### Statistical analysis

Analyses used standard machine-learning metrics implemented in scikit-learn. Binary discrimination was summarized by AUROC and average precision (reported in figures as PR-AUC). Threshold-dependent metrics included accuracy, balanced accuracy, precision, recall, specificity, and F1. Spearman and Pearson correlations were used for ranked or continuous outcomes as specified. Mann-Whitney U tests compared pathogenic and benign score distributions in supervised ClinVar analyses. CATH-Gene3D ^17^ and Reactome ^18^ nearest-neighbor analyses used cosine distance, and FlexiNR ^20^ feature families and context windows were compared using cross-validated metrics.

## Acknowledgements

We would like to thank all lab members in Wang Genomics Lab for the advice and suggestions. We appreciate the open-access UniRef and InterPro databases that make this study feasible. This project is supported by NIH grant HG013031 and the CHOP Research Institute

## Data Availability

The source resources used in this study-UniRef50 ^3^, InterPro ^2^, ClinVar ^4^, CATH-Gene3D ^17^, Reactome ^18^, PubMind ^19^, dbNSFP ^22^, MaveDB ^31^ and the published FlexiNR screen ^20^ are available from the repositories and publications cited here. The PALM model checkpoints and fine-tuned pathogenicity head are available for download at Hugging Face: https://huggingface.co/Wangwpi/PALM.

## Code Availability

The codes for the training, development and evaluation of PALM available at GitHub: https://github.com/WGLab/PALM.

## Supplementary Information

Supplementary materials contain extended methods, additional tables, and supplementary figures.

