## Supplementary materials for "A protein language model for domain, functional site, and variant effect prediction"

### Supplementary Figures

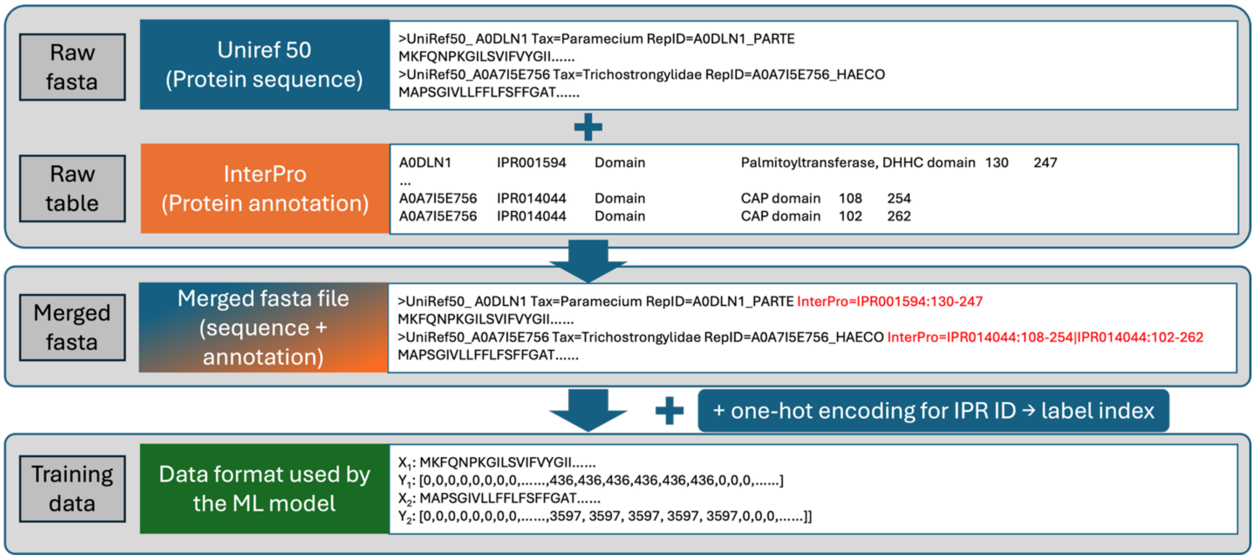


**Supplementary Figure 1. Data-processing workflow.** Raw inputs consist of protein sequences in FASTA format and InterPro annotations [1] in TSV format. InterPro identifiers and coordinate ranges are written to each FASTA header and subsequently converted into indexed residue-level labels for model training. For the model training using multi-label classification, the final assignment of label is not forced, the representation is a sets of all corrected labels instead of just one label.

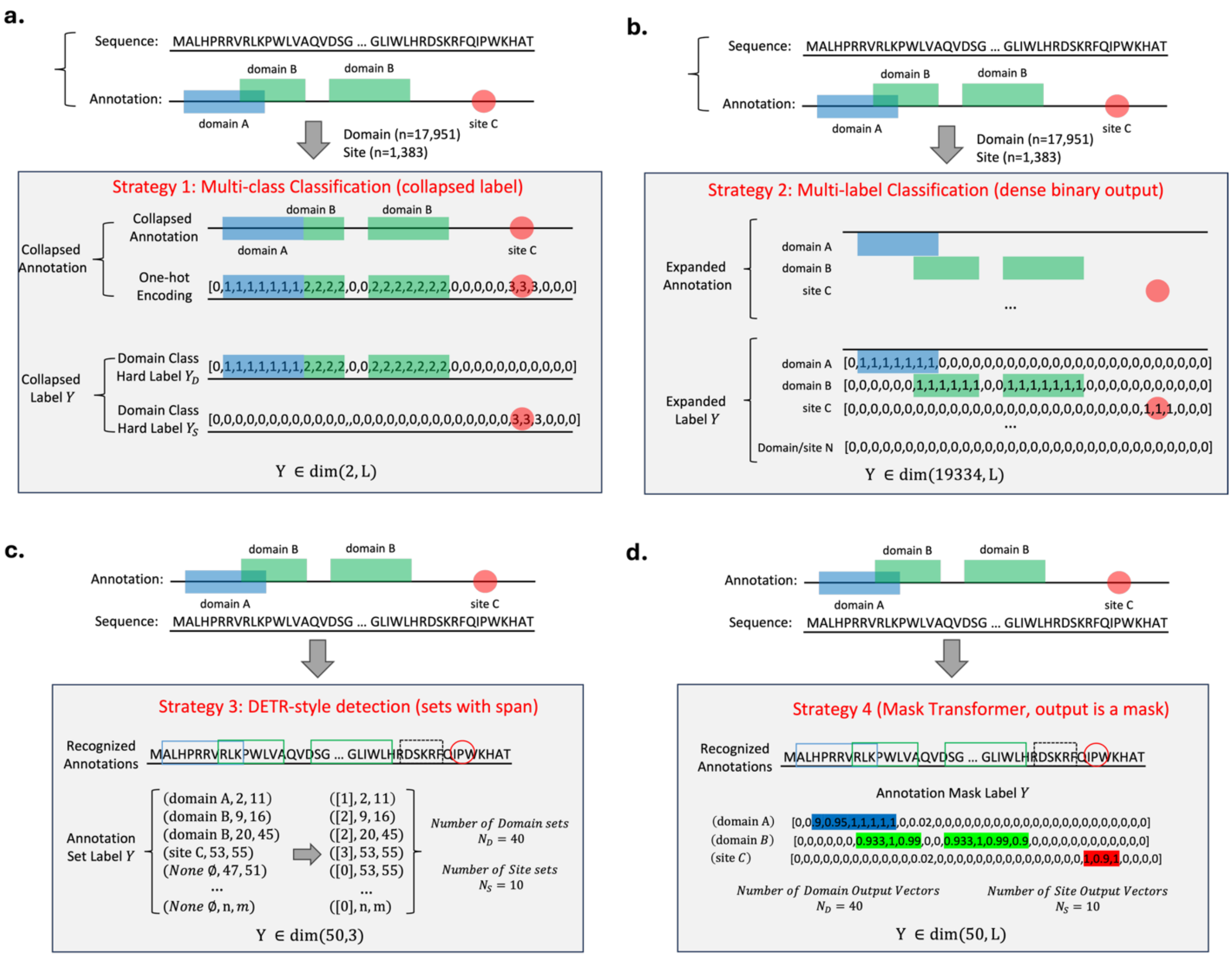


**Supplementary Figure 2. Four strategies for residue-level domain and site annotation.** Given a UniRef50 protein sequence [2] and its InterPro annotations [1], we evaluated four approaches to label construction and model training. **(a)** Strategy 1 treats each residue as a multiclass classification problem and retains only one final domain or site class. **(b)** Strategy 2 uses multilabel classification to preserve multiple annotations at each position, but requires a binary prediction for every possible class and therefore produces an output space dominated by zeros because annotations are sparse. **(c)** Strategy 3 formulates annotation as DETR-style set prediction, in which the model outputs a predefined number of sets containing a class, start position, and end position (implemented as center and width). **(d)** Strategy 4 uses a mask transformer to predict multiple annotation masks, allowing discontinuous blocks of the same annotation, such as domain B, to be represented jointly rather than as separate sets.

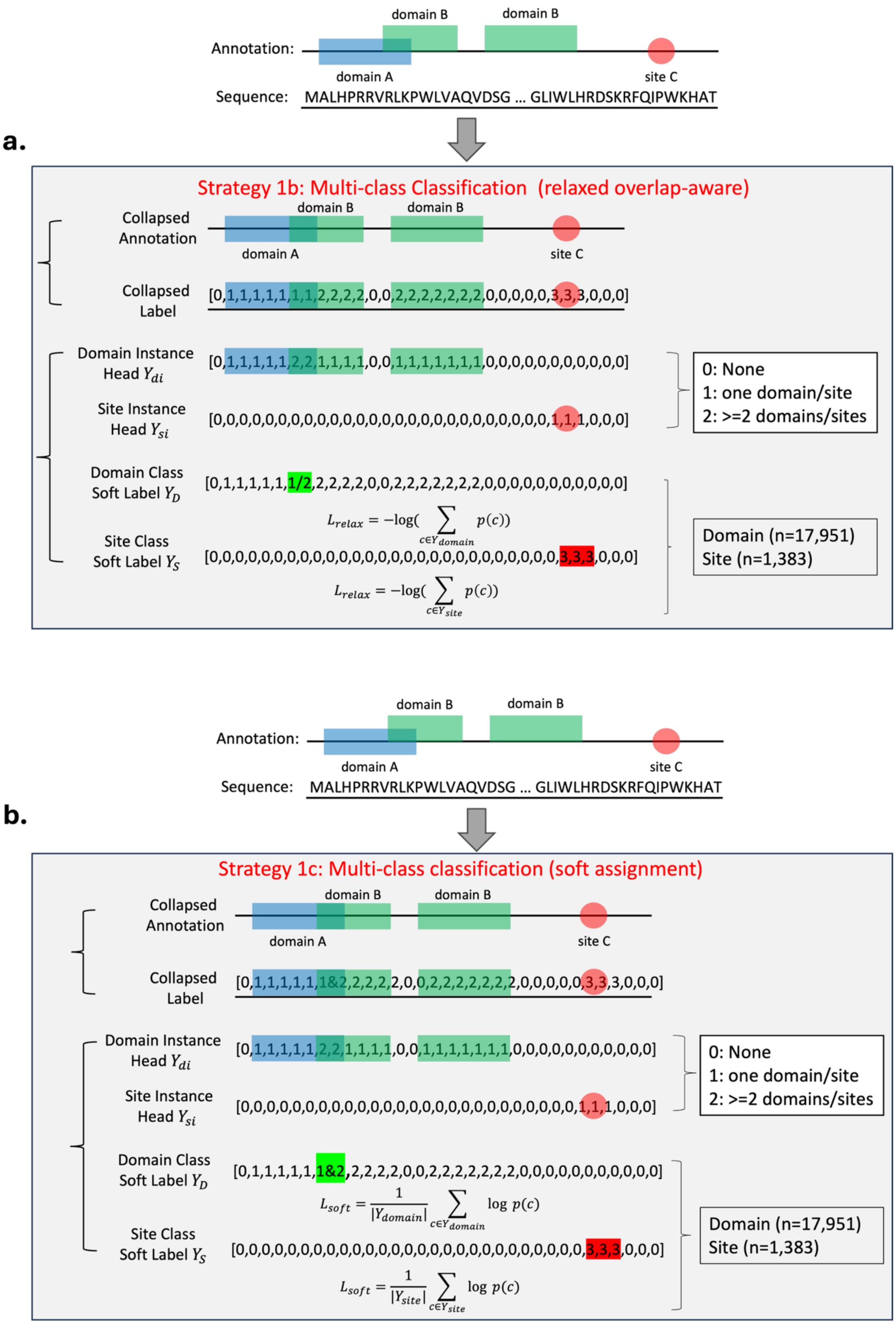

a.
Annotation:
domain B
domain B
domain A
site C
Sequence: MALHPRRVRLKPWLVAQVDSG ... GLIWLHRDSKRFQIPWKHAT
Strategy 1b: Multi-class Classification (relaxed overlap-aware)
Collapsed Annotation
Collapsed Label
[0,1,1,1,1,1,1,1,2,2,2,2,0,0,2,2,2,2,2,2,2,0,0,0,0,0,3,3,3,0,0,0]
Domain Instance Head $Y_{di}$
[0,1,1,1,1,1,2,2,1,1,1,1,0,0,1,1,1,1,1,1,1,0,0,0,0,0,0,0,0,0,0,0]
Site Instance Head $Y_{si}$
[0,0,0,0,0,0,0,0,0,0,0,0,0,0,0,0,0,0,0,0,0,0,0,0,0,0,1,1,1,0,0,0]
0: None
1: one domain/site
2: >=2 domains/sites
Domain Class Soft Label $Y_D$
[0,1,1,1,1,1,1/2,2,2,2,2,0,0,2,2,2,2,2,2,2,0,0,0,0,0,0,0,0,0,0,0]
$L_{relax} = -\log(\sum_{c \in Y_{domain}} p(c))$
Site Class Soft Label $Y_S$
[0,0,0,0,0,0,0,0,0,0,0,0,0,0,0,0,0,0,0,0,0,0,0,0,0,0,0,3,3,3,0,0,0]
$L_{relax} = -\log(\sum_{c \in Y_{site}} p(c))$
Domain (n=17,951)
Site (n=1,383)
b.
Annotation:
Sequence: MALHPRRVRLKPWLVAQVDSG ... GLIWLHRDSKRFQIPWKHAT
Strategy 1c: Multi-class classification (soft assignment)
Collapsed Annotation
Collapsed Label
[0,1,1,1,1,1,1&2,2,2,2,2,0,0,2,2,2,2,2,2,2,0,0,0,0,0,3,3,3,0,0,0]
Domain Instance Head $Y_{di}$
[0,1,1,1,1,1,2,2,1,1,1,1,0,0,1,1,1,1,1,1,1,0,0,0,0,0,0,0,0,0,0,0]
Site Instance Head $Y_{si}$
[0,0,0,0,0,0,0,0,0,0,0,0,0,0,0,0,0,0,0,0,0,0,0,0,0,0,1,1,1,0,0,0]
0: None
1: one domain/site
2: >=2 domains/sites
Domain Class Soft Label $Y_D$
[0,1,1,1,1,1,1&2,2,2,2,2,0,0,2,2,2,2,2,2,2,0,0,0,0,0,0,0,0,0,0,0]
$L_{soft} = \frac{1}{|Y_{domain}|} \sum_{c \in Y_{domain}} \log p(c)$
Site Class Soft Label $Y_S$
[0,0,0,0,0,0,0,0,0,0,0,0,0,0,0,0,0,0,0,0,0,0,0,0,0,0,0,3,3,3,0,0,0]
$L_{soft} = \frac{1}{|Y_{site}|} \sum_{c \in Y_{site}} \log p(c)$
Domain (n=17,951)
Site (n=1,383)


23

**Supplementary Figure 3. Adaptations of the multiclass strategy for overlap-aware training. (a)** In the relaxed objective, the probabilities assigned to valid classes are summed before loss calculation; the model is therefore not penalized for concentrating on one correct class when multiple classes are valid. **(b)** PALM uses a soft-target multiclass objective. InterPro annotations [1] are collapsed while retaining overlapping labels, and target probability is distributed across all valid domain or site classes. This objective encourages recovery of multiple annotations when they overlap at a residue.

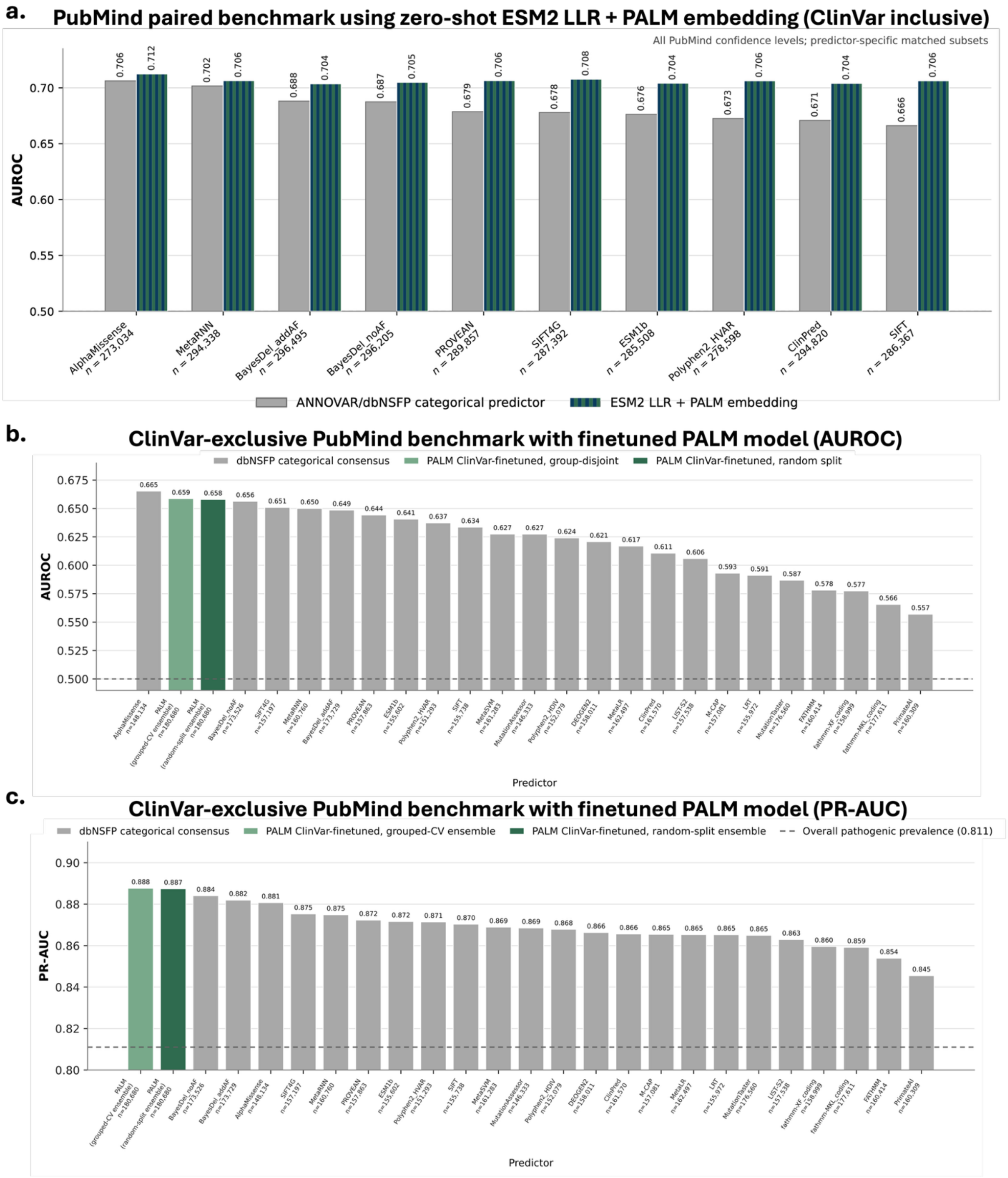


**Supplementary Figure 4. Zero-shot and ClinVar [3] fine-tuned PALM evaluation on PubMind [4].**

**(a)** Pairwise comparison of the combined zero-shot ESM2 LLR–PALM embedding score with the ten highest-ranked ANNOVAR[5]/ dbNSFP [6] categorical predictors across all PubMind confidence levels (including variants overlapped with ClinVar). The total number of variants in each pairwise comparison is shown in the plot. **(b, c)** AUROC (b) and PR-AUC (c) for two five-fold ensembles of ClinVar-fine-tuned pathogenicity heads, trained using random-stratified or gene/protein-disjoint splits, compared with 23 ANNOVAR/dbNSFP [5, 6] predictors. ClinVar-exclusive PubMind varaints are used in the comparison. PALM ensemble scores are unweighted means of the five fold-level pathogenicity probabilities. The dashed line in (c) indicates the PubMind pathogenic prevalence (0.811). Values above the bars report AUROC or PR-AUC.

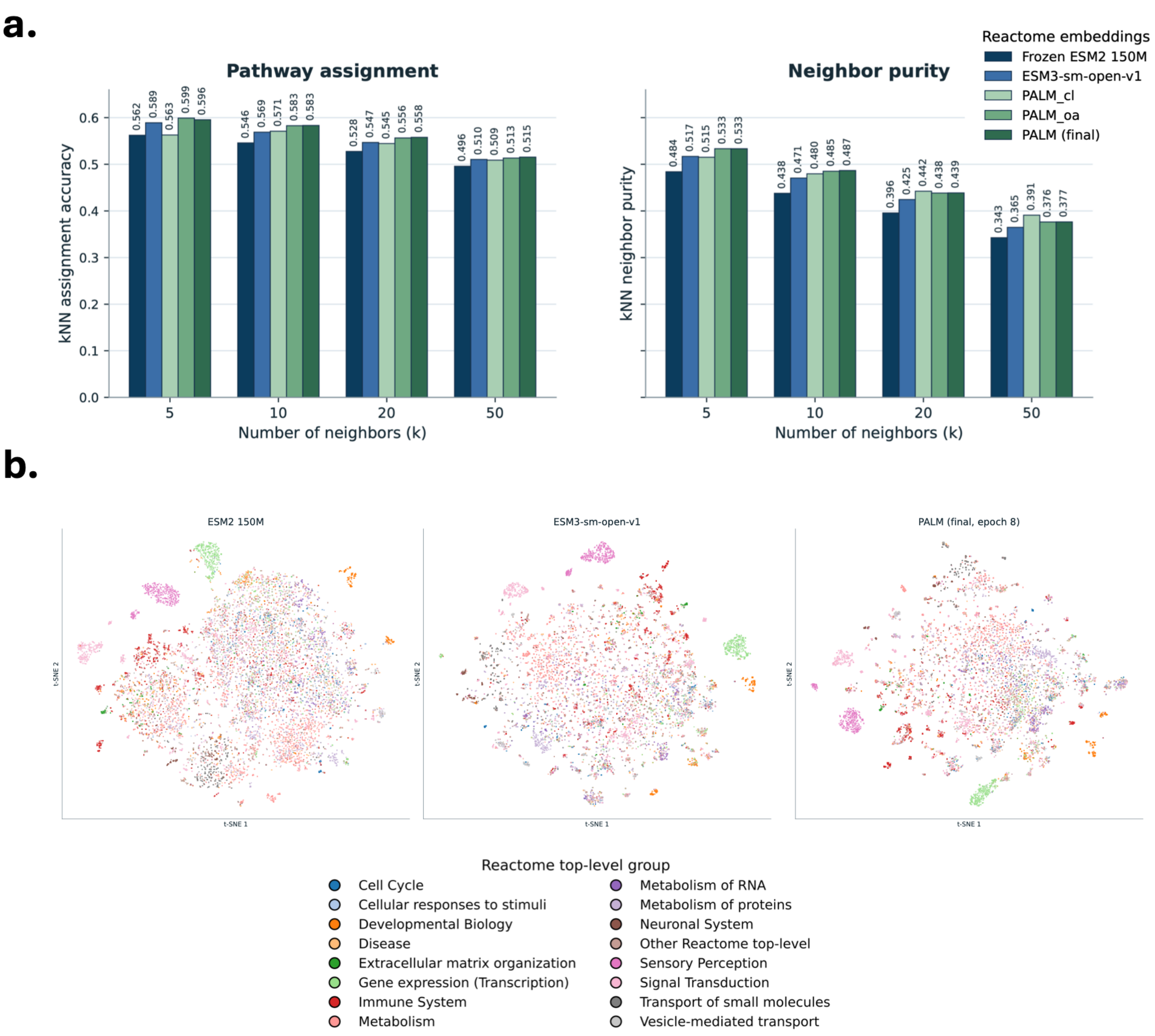


**Supplementary Figure 5. PALM representations capture Reactome pathway-level organization [7]. (a)** Reactome pathway-assignment accuracy and neighbor purity for frozen ESM2-150M [8], ESM3-sm-open-v1 [9], and PALM embeddings across k = 5-50 nearest neighbors (11,312 proteins; 16 pathway labels). **(b)** t-SNE visualization of ESM2, ESM3, and final PALM embeddings colored by the 16 top-level Reactome pathway groups. PALM shows clearer pathway-associated separation and grouping than the pretrained ESM baselines in this complementary functional benchmark.

### Supplementary Tables

**Supplementary Table 1**. Held-out test set stratified by protein sequence-identity (PID).

| **Maximum PID to train/validation** | **Test proteins, n** | **PID subset** | **Full test set** |
|---|---|---|---|
| <20% or no detectable hit | 25,149 | 3.66% | 1.09% |
| 20–<30% | 44,410 | 6.47% | 1.92% |
| 30–<40% | 107,555 | 15.67% | 4.66% |
| 40–<50% | 195,524 | 28.49% | 8.47% |
| 50–<70% | 183,087 | 26.67% | 7.93% |
| 70–<90% | 94,376 | 13.75% | 4.09% |
| 90–100% | 36,269 | 5.28% | 1.57% |
| **Total PID subset** | **686,370** | **100.00%** | **29.72%** |

**Supplementary Table 2. Performance of different model architectures and settings.** An asterisk indicates that all InterPro site types were included, including repeats and post-translational modification (PTM) sites.

| **Model** | **Best Epoch (Total)** | **Time for 1 epoch** | **Domain Type Accuracy** | **Site Presence Precision** | **Site Presence Recall** | **Site Presence F1** | **Site Type Accuracy** |
|---|---|---|---|---|---|---|---|
| BiLSTM$_{8M}$ | 9 (10) | 399.8 min | 0.9050 | 0.525 | 0.878 | 0.657 | 0.9922 |
| BiLSTM$_{150M}$ | 6 (6) | 1058.7 min | 0.9345 | **0.619** | 0.950 | **0.750** | **0.9937** |
| BiLSTM$_{650M}$ | 2 (2) | 1710.8 min | **0.9362** | 0.529 | **0.970** | 0.685 | 0.9929 |
| Transformer$_{8M}$ | 13 (16) | 362.5 min | 0.8735 | 0.375 | 0.910 | 0.531 | 0.9884 |
| LoRA$_{150M}$*+MLP | 6(6) | 1057.5 min | 0.9138 | 0.404 | 0.870 | 0.552 | 0.9913 |
| BiLSTM$_{650M}$* | 2(2) | 1736.5 min | 0.9343 | 0.532 | 0.966 | 0.686 | 0.9925 |

| | | | | | | | |
|---|---|---|---|---|---|---|---|
| LoRA+BiLSTM$_{150M}$ (subset)* | 10(10) | 96.7 min | 0.9024 | 0.621 | 0.920 | 0.741 | 0.9815 |
| LoRA+BiLSTM$_{150M}$ * | 7(9) | 760.6 min | 0.9402 | 0.751 | 0.962 | 0.843 | 0.9955 |
| LoRA+BiLSTM$_{650M}$ * | 6(6) | 1036.0 min | 0.9458 | 0.753 | 0.972 | 0.849 | 0.9953 |
